# Electron tomography reveals mitochondrial network and cristae remodelling during cell differentiation in the human placenta

**DOI:** 10.1101/2025.06.09.658204

**Authors:** Siddharth Acharya, Eric Hanssen, Veronica B. Botha, Tia M. Smith, Sahan Jayatissa, Zlatan Trifunovic, Lucy A. Bartho, John E. Schjenken, Tu’uhevaha J. Kaitu’u-Lino, Anthony V. Perkins, Joanna L. James, Kirsty G. Pringle, James C. Bouwer, Roger Smith, Joshua J. Fisher

**Affiliations:** School of Biomedical Sciences and Pharmacy, College of Health, Medicine and Wellbeing, University of Newcastle, Callaghan, NSW, Australia; Ian Holmes Imaging Centre and ARC Industrial Training Centre for Cryo-electron Microscopy of Membrane Proteins, The Bio21 Molecular Science and Biotechnology Institute, University of Melbourne, Parkville, VIC, Australia; Department of Biochemistry and Pharmacology, University of Melbourne, Parkville, VIC, Australia; School of Environmental and Life Sciences, College of Engineering, Science and Environment, University of Newcastle, Callaghan, NSW, Australia; Auckland Bioengineering Institute, University of Auckland, Auckland, New Zealand; Translational Obstetrics Group, The Department of Obstetrics, Gynaecology and Newborn Health, Mercy Hospital for Women, University of Melbourne, Heidelberg, VIC, Australia; Infertility and Reproduction Research Program, Hunter Medical Research Institute, New Lambton Heights, NSW, Australia; School of Health, University of the Sunshine Coast, Sippy Downs, QLD, Australia; Department of Obstetrics, Gynaecology and Reproductive Sciences, Faculty of Medical and Health Sciences, University of Auckland, Auckland, New Zealand; Women’s Health Research Program, Hunter Medical Research Institute, New Lambton Heights, NSW, Australia; Molecular Horizons and School of Chemistry and Molecular Bioscience, University of Wollongong, NSW, Australia; ARC Industrial Transformation Training Centre for Cryo-electron Microscopy of Membrane Proteins, University of Wollongong, NSW, Australia; School of Medicine and Public Health, College of Health, Medicine and Wellbeing, University of Newcastle, Callaghan, NSW, Australia; Mothers and Babies Research Program, Hunter Medical Research Institute, New Lambton Heights, NSW, Australia

**Author notes:** Corresponding Author*: Dr. Joshua J. Fisher, Postdoctoral Research Fellow, Mothers and Babies Research Program, Hunter Medical Research Institute, University of Newcastle.

**Keywords:** array tomography, volume imaging, cryo-electron tomography, mitochondrial dynamics, cristae architecture, electron transport chain, supercomplex, cellular differentiation, trophoblast, placenta

## Abstract

Mitochondria adapt their structure through fusion and fission, yet how their morphology and cristae architecture alter as cells differentiate remains unclear. The human placenta is an ideal model for studying mitochondrial diversity within a single tissue. As epithelial trophoblast cells of the placenta differentiate, their accompanying mitochondria morphologically and functionally transform. We used array tomography to characterise mitochondrial volume and network complexity. Cryo-electron tomography revealed two distinct subpopulations of mitochondria within progenitor cytotrophoblasts, and a singular, homogeneous population in the differentiated syncytiotrophoblast. We showcased the 3D topology of individual cristae through a standardised metric of “curvedness”. Proteomic analysis of isolated mitochondria identified reduced dynamics proteins and cristae organisation complexes, accompanied by subunit variations within the electron transport chain, ATP synthase, and altered supercomplex abundance. This study highlights the advantages of combining multimodal imaging with mechanistic insights, to elucidate the process of mitochondrial morphology and cristae architecture remodelling as cells differentiate.

## Introduction

Mitochondria are intracellular organelles responsible for energy production, steroid hormone synthesis, and the regulation of cellular metabolism and immunity.^1^ To support the bioenergetic demands of the cell, mitochondria produce energy in the form of adenosine triphosphate (ATP), which facilitates cellular proliferation, migration, and the catalysis of biochemical reactions.^1^ Mitochondria adapt their bioenergetic capacity and respiratory rate to support cellular demand, by remodelling their gross network structure through a continuous process of fusion and fission.^2^ Concurrently, the remodelling of internal mitochondrial architecture, the intermembrane space (area between the outer and inner mitochondrial membranes), and folding of the inner membrane into cristae, enables each mitochondrion to adapt for either high bioenergetic demands or steroidogenic roles.^3^ Despite the significance of mitochondrial structure and its role in dictating functional capacity, our understanding of the underlying protein mechanisms remains limited.

Mitochondrial fusion occurs when cellular demands for ATP are high, resulting in the formation of interconnected structures with an increased surface area and volume for efficient ATP synthesis. This is achieved by two proteins; mitofusin 2 (MFN2) that enables the fusion of the mitochondrial outer membrane, and optic atrophy 1 (OPA1) that promotes the fusion of the inner membrane. Together, these proteins ensure that fused mitochondria maintain a continuous internal structure for the distribution of mitochondrial DNA, proteins, and metabolites.^2,4,5^ Fused mitochondrial networks are common in human tissues such as skeletal muscle, which are characterised by high bioenergetic demands.^6^ Conversely, when cellular demands are reduced, mitochondrial fission leads to the fragmentation of mitochondrial networks, an adaptation that supports basal ATP synthesis and specialised roles such as steroidogenesis, mitochondrial biogenesis, and the clearance of dysfunctional mitochondria (mitophagy) during cellular senescence.^2,7–10^ Fission is driven by the presence of mitochondrial fission protein 1 (FIS1), which recruits dynamin-related protein 1 (DRP1) for cleavage.^4,11^ DRP1 proteins bind to FIS1 receptors on the surface of the mitochondrial outer membrane, and form spiral-like structures that constrict and separate mitochondria into separate organelles.^12^ In certain cell types such as the early germ cells of the testes, fragmented mitochondria have been shown to be physiologically beneficial, exhibiting a protective effect against the accumulation of reactive oxygen species.^13^ Fission events also facilitate the distribution of mitochondria to distal regions of cells with specific functions, as observed in neuronal axons.^14^

The internal architecture of mitochondria is critical for regulating bioenergetic function and the efficiency of ATP synthesis. The organisation of the inner membrane and formation of cristae is controlled by the mitochondrial intermembrane space bridging (MIB) complex.^15^ The MIB complex comprises several protein complexes within the outer and inner mitochondrial membranes, most notably the mitochondrial contact site and cristae organising system (MICOS) proteins. The MICOS complex, consisting of at least seven different proteins, is located within the inner membrane, and is suggested to associate and interact with three proteins within the sorting and assembly machinery (SAM) complex and at least two metaxin (MTX) proteins of the outer membrane.^15–17^ The interaction between MIB complexes is critical for the organisation and structure of cristae, the formation of the intermembrane space, and the stability of inner membrane curvature.^15^ The resulting diverse range of cristae architectures reflects cellular and tissue demands and functions. Conventionally, mitochondria within cardiac and skeletal muscle tissue contain stacked lamellar cristae,^18–21^ with the majority of the matrix (inner cavity of the mitochondrion) occupied by closely packed, narrow folds with constricted crista junctions.^22,23^ In contrast, steroidogenic tissues such as the adrenal cortex and testis have mitochondria with rounded cristae, optimising the structure of the inner membrane for the import of cholesterol and subsequent synthesis of steroid hormones.^24,25^ Recent advances in electron microscopy have revealed a diverse range of cristae architectures, including tubular, vesicular, onion, or any combination of these – each with contrasting volumetric properties that enable diverse bioenergetic functions.^26,27^ The exact mechanism of interaction between MIB proteins and the formation of diverse cristae structures remains unclear,^28^ although loss of MICOS results in overall MIB disassembly, and the formation of unfolded globular and spherical cristae, demonstrating its critical role.^29,30^ The subsequent loss of cristae and inner membrane surface area, is thought to limit the efficiency of the electron transport chain (ETC), a series of protein complexes that are critical for ATP synthesis.^31^

The synthesis of ATP is supported by the ETC, which is comprised of complex I (NADH dehydrogenase), complex II (succinate dehydrogenase), complex III (Q-cytochrome c oxidoreductase), and complex IV (cytochrome c oxidase). Collectively, the complexes in mammalian mitochondria consist of at least 73 subunit proteins, and are typically categorised based on their function. Core subunits provide functional redox reaction centres for the transport of electrons, while accessory subunits provide structural stability and enable conformational changes to occur that assist in proton translocation.^32,33^ As electrons move through complexes I, III, and IV, these conformational shifts pump protons into the intermembrane space. As a result, an electrochemical gradient is formed across the inner mitochondrial membrane.^34–39^

To equilibrate the electrochemical gradient, protons must move across the inner membrane towards the relatively low proton concentration of the mitochondrial matrix.^40,41^ While the inner membrane itself is not permeable to protons, a channel for proton translocation is provided by ATP synthase.^42^ The movement of protons through ATP synthase provides sufficient kinetic energy for the phosphorylation of adenosine diphosphate into ATP.^43^ Studies have identified the capacity of ATP synthase to oligomerise, whereby several ATP synthase units assemble together to optimise bioenergetic efficiency and improve structural stability.^44,45^ In human kidney cells, ATP synthase dimers have been observed to line mitochondrial cristae, inducing curvature of the inner membrane to form sharp cristae apices and topologically complex ridges.^46–49^ Lamellar cristae apices in particular, form marked invaginations that shape the inner membrane into a funnel-like pocket, where protons rapidly move towards ATP synthase dimers located at the apex, thus increasing the rate and efficiency of ATP synthesis.^47,49–51^ Lack of ATP synthase dimers leads to the formation of atypical onion and globular cristae.^52,53^ Emerging research proposes that specific subunits such as ATP5ME, ATP5MG, and ATP5MK have a critical role in coordinating the dimerisation of ATP synthase in yeast mitochondria.^49^ Similarly, accessory subunits within complexes I, III, and IV have been suggested to play a role in the assembly of multiple complexes into higher-order structures known as supercomplexes.^54^

Supercomplexes are highly efficient assemblies that supersede the kinetic and functional limits of individual complexes. Supercomplexes have been documented to assemble in the forms I+III+IV or III+IV, and may coexist simultaneously with individual complexes in a dynamic state of plasticity.^55^ Studies report the presence of supercomplexes when cellular bioenergetic demands are high, increasing the efficiency of ATP synthesis by forming tunnels for steady electron flux, instead of undergoing sequential complex-to-complex electron transport, which is rate-limited.^56,57^ Furthermore, supercomplexes mitigate the leakage of electrons by eliminating the need for electron shuttling, and by providing a surrounding subunit structure that protectively entraps electrons within the assembly. As a result, supercomplexes limit the formation of reactive oxygen species – a toxic byproduct caused by the direct reaction of electrons with oxygen molecules in the mitochondrial matrix, forming free radicals that inflict cellular damage.^58^ While the presence of ATP synthase dimers and supercomplexes have primarily been characterised in plant and mammalian species, their role in human cell biology is not fully understood. There is limited consensus on whether membrane curvature is solely induced by the presence of ATP synthase oligomers and supercomplexes, or conversely, whether curved membranes stabilise and promote the formation of higher-order complex structures.^59,60^

Our understanding of the interplay between mitochondrial dynamics, cristae architecture, the incorporation of ETC complexes, and how the assembly of ATP synthase dimers and supercomplexes converge to remodel mitochondrial structure remains incomplete. Defining the orientation and complex topology of mitochondria and internal cristae requires 3D electron microscopy techniques to adequately assess functionally significant characteristics of surface area and volume. The diversity of mitochondrial structure has been the subject of investigations in multiple species, including human cerebral, cardiac, and hepatic tissues.^19,61,62^ We propose that the placenta is an ideal model for studying mitochondrial structure within a singular tissue environment as it exhibits dramatic differences in mitochondrial morphology that can be visualised within neighbouring progenitor and differentiated cell populations. The unique architecture of the placenta supports the diverse functions required to maintain pregnancy including maternal-fetal exchange, steroidogenesis, and the synthesis of signalling molecules that regulate maternal physiology.^63^ The placenta achieves this via its branched structure, in which fetal-derived placental villi are bathed in maternal blood to permit the exchange of nutrients, gases, and wastes between maternal and fetal circulations.^64–66^ Villi are lined with a trophoblast bilayer, which consists of inner cytotrophoblast (CTB) cells that fuse together and terminally differentiate into the outer, multinucleated syncytiotrophoblast (STB).^64^ In contrast to the proliferative CTB, the STB has been shown to have a lower capacity for producing ATP, an adaptation thought to be driven by its specialised roles in mediating biochemical reactions and synthesising steroid hormones critical for pregnancy.^67–69^

Using transmission electron microscopy (TEM), it was first observed in the 1990’s that mitochondria within neighbouring cell populations of the placenta have diverse morphologies, in which mitochondria of CTB have a larger 2D area in comparison to compact mitochondria of the adjacent STB.^69^ Martinez et al.,^69^ proposed that the greater surface area to volume ratio of mitochondria within the STB may enable the efficient import of cholesterol and subsequent synthesis of pregnenolone. Subsequent studies have shown that the STB has a lower bioenergetic demand than the CTB, supported by reduced mitochondrial functionality.^67,70,71^ While CTBs and the STB have varied bioenergetic demands, the ways in which mitochondrial network structure and cristae architecture remodel in response to trophoblast differentiation, remain unclear. Such a characterisation demands the application of 3D electron microscopy to study intact mitochondrial networks within placental tissue, and a high-resolution approach that can reveal the internal architecture of placental mitochondria. While 3D electron microscopy has previously been applied to study the structure of the trophoblast bilayer and mitochondria within the CTB,^72^ a comprehensive volumetric analysis of mitochondria from both CTB and STB populations does not currently exist.

We leveraged the unique architecture of the placenta, in which mitochondria from progenitor and differentiated cell populations exist in close proximity yet exhibit distinct structural and functional characteristics to support their contrasting bioenergetic requirements. We adopted a multimodal imaging approach, using array tomography and cryo-electron tomography (cryo-ET) to assess the complex 3D structure of mitochondrial networks and topology and the curvature of cristae architecture. Additionally, we characterised proteins involved in mitochondrial dynamics, the MIB complex, subunits of complexes I-IV and ATP synthase, and the presence of supercomplexes in the human placenta. Together, this provides new insights into the mechanisms underpinning the remodelling of mitochondrial structure, and how mitochondria adapt to support shifting cellular demands.

## Results

### Array tomography reveals the differences in mitochondrial network complexity between CTBs and the STB

To characterise the 3D structure of whole mitochondrial networks within CTBs (Cyto-Mitos) or the STB (Syncytio-Mitos), we performed array tomography on resin-embedded term placental villous tissue (Fig. 1A). Our reconstruction of the whole trophoblast bilayer (adjacent CTB and STB; Fig. 1B-D) showed the epithelial-like structure of the CTB, consisting of projections that surround the bordering basement membrane. Notably, despite being 0.6 µm apart, Cyto- and Syncytio-Mitos had contrasting network structures and morphologies. While Cyto-Mitos (pink) formed clusters and had a higher degree of interconnectedness, as reflected in their increased mitochondrial complexity index, Syncytio-Mitos (yellow) were fragmented and isolated (Fig. 1B-D, L). In addition to demonstrating contrasting gross network structures, our approach of reconstructing individual mitochondria built upon previously published 2D analyses,^67,69^ revealing a spectrum of morphologies including branched, elongated, and tubular mitochondria in the CTB, and small, spherical, and compact mitochondria in the STB (Fig. 1E, F). Cyto-Mitos were significantly larger and more elongated, with a mean surface area of 1.71 µm^2^ and volume of 0.135 µm^3^, compared to the smaller and more spherical Syncytio-Mitos, which had a mean surface area of 0.491 µm^2^ and volume of 0.0307 µm^3^ (Fig. 1G, H, J, K). Consequently, Cyto-Mitos had a lower surface area to volume ratio (Fig. 1I).

**Figure 1:**
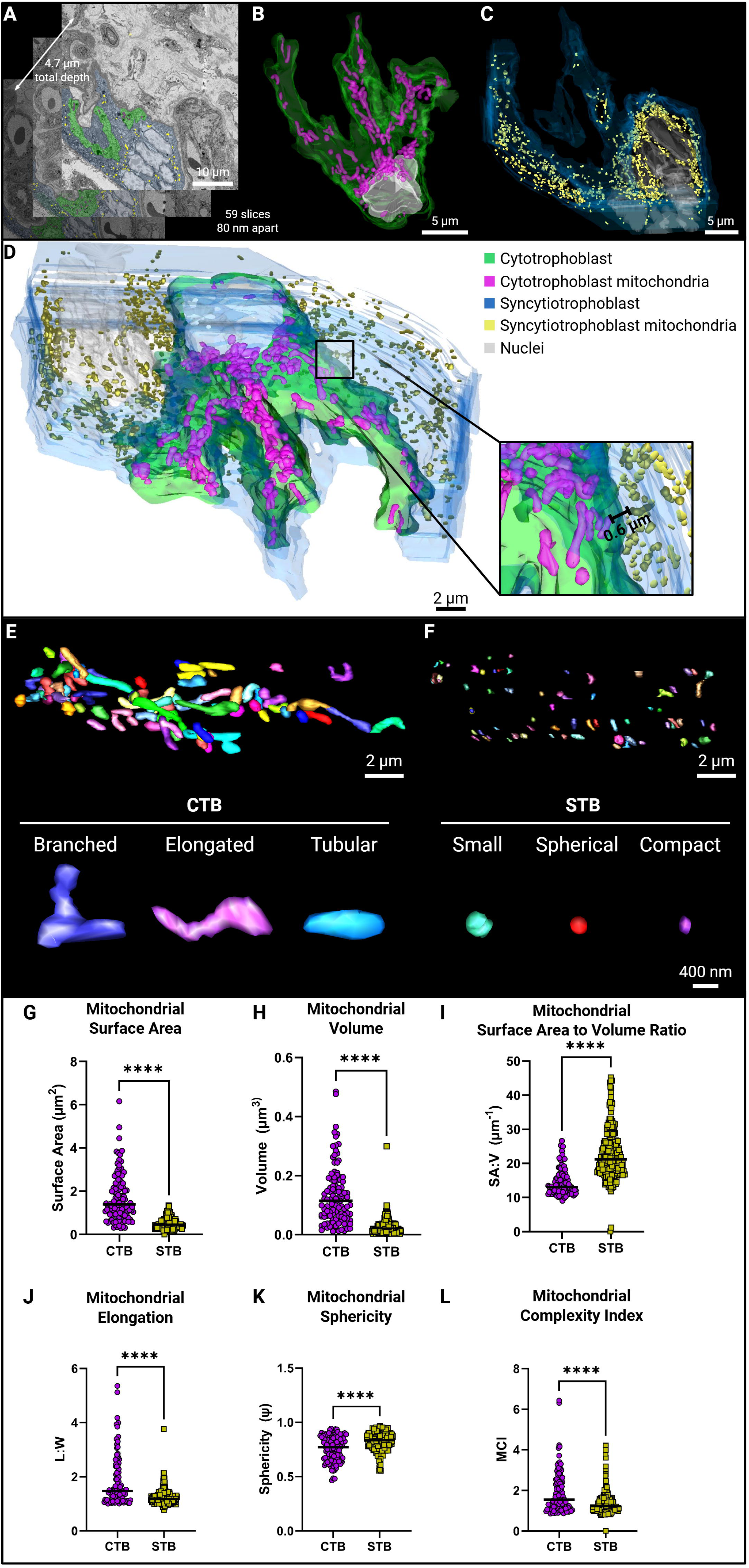
Array tomography of placental villi reveals mitochondrial networks of the CTB and STB. A) Representative Z-slices from our array tomography image series with a Z-depth of 4.7 µm, the full image series had a total Z-depth of 20.16 µm. B) 3D reconstruction of a CTB cell (green), nucleus (white), and CTB mitochondria (pink). C) 3D reconstruction of the STB (blue), nucleus (white), and STB mitochondria (yellow). D) Full in situ 3D reconstruction of a CTB and surrounding STB, revealing sub-micron distances between diverse mitochondrial structures. E) Randomly selected subset of mitochondria from the CTB and (F) STB, with representative branched, elongated, and tubular mitochondria from the CTB and small, spherical, and compact mitochondria of the STB. G-L) Volumetric analysis was performed on randomly selected subsets of mitochondria obtained from two CTBs (n = 113 Cyto-Mitos) and two STBs (n = 241 Syncytio-Mitos), collectively obtained from two healthy full-term placentae. The graphs presented are G) surface area, H) volume, I) surface area to volume ratio, J) elongation, K) sphericity, and L) mitochondrial complexity index. Mann-Whitney U tests were performed to examine significant variations between groups, expressed as p < 0.05*, p ≤ 0.01**, p ≤ 0.001***, p ≤ 0.0001****.

### Transformation of mitochondrial cristae architecture following trophoblast differentiation

Whilst array tomography enabled the observation of intact mitochondrial networks within their native tissue environment, its resolution and scale limited the investigation of internal mitochondrial membranes such as cristae, which serve as key structures that house the functional units of mitochondrial bioenergetics. Therefore, we used cryo-ET to increase the imaging resolution ∼ 260-fold, to focus on imaging each individual mitochondrion isolated from CTBs and the STB as previously described.^67^ Following tomogram acquisition and 3D reconstruction of internal mitochondrial architecture (Fig. 2A), we observed a greater abundance of invaginated cristae in Cyto-Mitos, compared with Syncytio-Mitos (Fig. 2B-D). For the first time, we observed two distinct mitochondrial subpopulations in Cyto-Mitos; those with lamellar cristae, formed by long inner membrane folds and sharp apex curvatures,^73^ and those with tubulo-vesicular cristae, which exhibited more rounded and bulbous structures.^23^ In contrast, Syncytio-Mitos only exhibited globular cristae architecture, characterised by rounded and spherical structures with no drastic deviations in curvature.^74^

**Figure 2:**
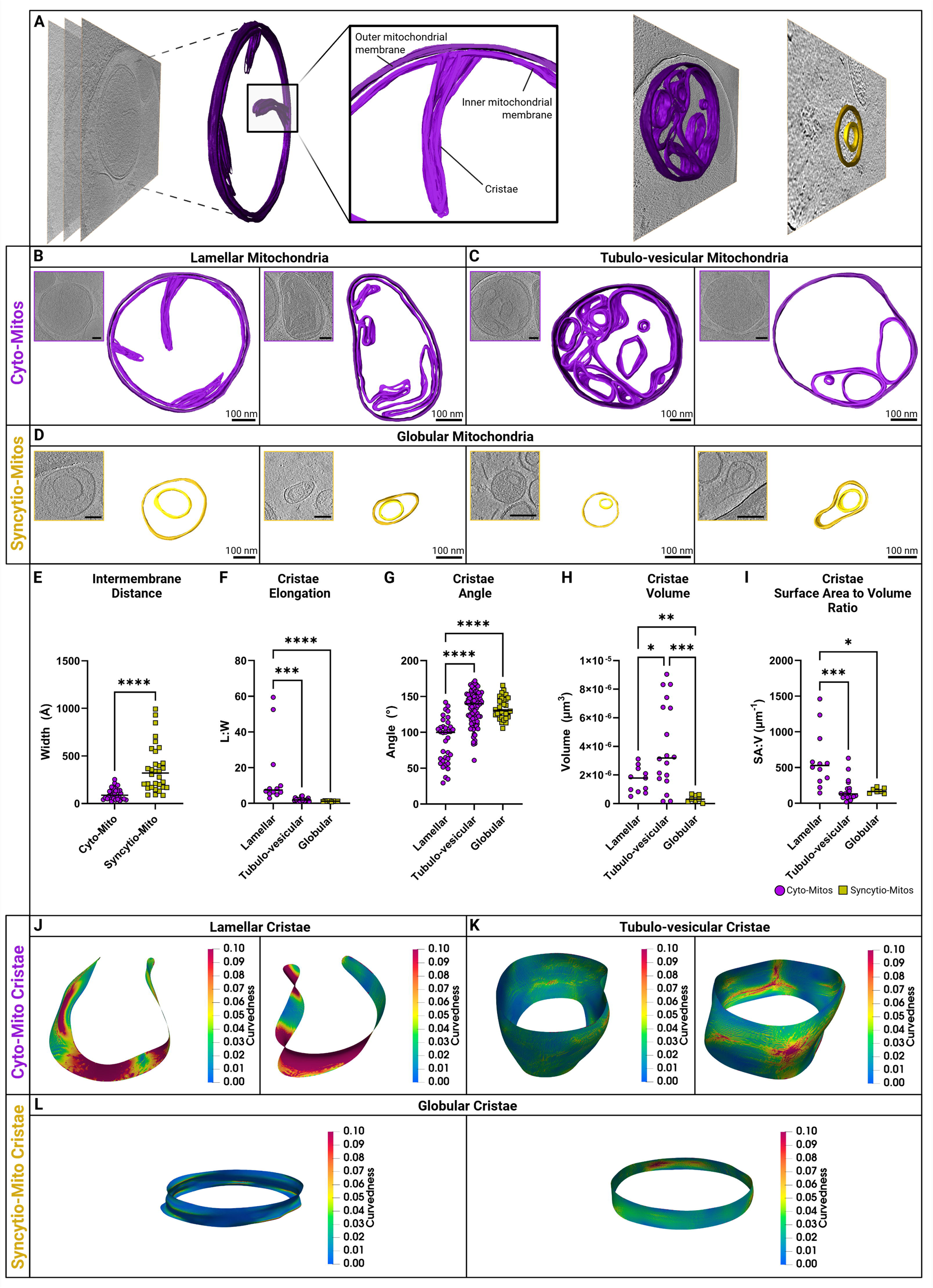
Cryo-ET analysis of internal mitochondrial architecture within Cyto- and Syncytio-Mitos. A) 16 mitochondria from two healthy full-term placentae were imaged, with representative tomographic slices of Cyto-Mitos (purple) and Syncytio-Mitos (yellow) shown. B) Representative Cyto-Mitos that consist of lamellar cristae and C) tubulo-vesicular cristae, and D) representative Syncytio-Mitos that consist of globular cristae. All scalebars represent 100 nm. E-I) Volumetric analyses were performed to quantify and compare cristae architecture between Cyto-Mitos (purple circles) and Syncytio-Mitos (yellow squares). E) Intermembrane distance was compared between Cyto-Mitos and Syncytio-Mitos where each data point represents an independent width measurement between the outer and inner mitochondrial membranes (n = 31 measurements for Cyto-Mitos, n = 32 measurements for Syncytio-Mitos). F) Cristae elongation was assessed for each lamellar (n = 12 cristae), tubulo-vesicular (n = 19 cristae), and globular cristae (n = 8 cristae), between Cyto- and Syncytio-Mitos, where each data point represents an individual cristae. G) Cristae angle was measured for lamellar (n = 36 measurements), tubulo-vesicular (n = 85 measurements), and globular (n = 32 measurements) cristae. Each data point represents an independent angle measurement of functionally pertinent zones of the cristae (apices and junctions for lamellar cristae, or using a quadrant system for tubulo-vesicular and globular cristae). Additionally, H) cristae volume and I) surface area to volume ratio was calculated using 3D reconstructions. Each data point represents an individual lamellar (n = 12 cristae), tubulo-vesicular (n = 19 cristae), and globular (n = 8 cristae) cristae. J-L) 3D curvature was assessed across six cristae reconstructions, using the curvedness formula. Intermembrane distance was assessed using the Mann-Whitney t test, while volume was assessed using the Brown-Forsythe and Welch ANOVA test (post-hoc). All other cristae metrics were assessed using the Kruskal-Wallis and Dunn’s multiple comparisons tests (post-hoc). Significant variation is expressed as p < 0.05*, p ≤ 0.01**, p ≤ 0.001***, or p ≤ 0.0001****.

Volumetric analyses of the cryo-ET data concluded that Cyto-Mitos had a significantly smaller intermembrane distance than Syncytio-Mitos (Fig. 2E). Our analysis of individual cristae architectures concluded that lamellar cristae were significantly more elongated (Fig. 2F) and had a more acute degree of curvature at their functionally pertinent zones (apices and junctions) than either tubulo-vesicular or globular cristae (Fig. 2G). Within Cyto-Mitos, lamellar cristae had a lower volume than tubulo-vesicular cristae. However, both architectures had a greater volume compared to the smaller globular cristae of Syncytio-Mitos (Fig. 2H). Subsequently, we assessed the functionally critical metric of surface area to volume ratio and determined that lamellar cristae of Cyto-Mitos had a significantly higher ratio than both tubulo-vesicular cristae and globular cristae (Fig. 2I). Building upon our 2D measurements, we characterised the complex topology and curvature of cristae by assessing the curvedness of the 3D cristae reconstructions, confirming that lamellar cristae had high curvedness localised to apices (Fig. 2J). Tubulo-vesicular cristae of Cyto-Mitos had uniformly distributed curvedness with localised pockets of angularity, while globular cristae of Syncytio-Mitos had the lowest curvedness compared to both subpopulations of Cyto-Mitos (Fig. 2K, L).

### Reduction in cristae organisation protein levels underpins the structural remodelling of syncytialised mitochondria

To understand the mechanisms underpinning the dramatic alterations in mitochondrial structure observed between Cyto- and Syncytio-Mitos following differentiation, we isolated each mitochondrial population from fresh placental tissue and performed an in silico analysis of liquid chromatography-mass spectrometry (LC-MS) data. This enabled the examination of mitochondria-specific proteins that govern cristae architecture. When investigating the abundance of proteins involved in mitochondrial dynamics (the balanced interplay of fusion and fission), our findings indicated an increase of fusion proteins MFN2 (0.64 log_2_ fold change (log_2_FC)) and OPA1 (0.16 log_2_FC), and fission protein FIS1 (0.58 log_2_FC) in Cyto-Mitos. Additionally, there was a higher abundance of the fragmentation and cleavage protein DRP1 (-0.12 log_2_FC) in Syncytio-Mitos (Fig. 3A, B). Fragmentation of Syncytio-Mitos was confirmed by array tomography (Fig. 3C). Additionally, we investigated proteins of the MIB complex, including those within the SAM and MICOS complexes (Fig. 3D, E). We identified that proteins involved in connecting and tightening cristae junctions (including MTX2 (0.45 log_2_FC), MTX1 (0.74 log_2_FC), and SAMM50 (0.81 log_2_FC)) were more abundant in Cyto-Mitos, whereas the inner membrane stability protein DNAJC11 (-0.21 log_2_FC) was more abundant in Syncytio-Mitos. Within the MICOS complex, which has a critical role in maintaining the folding of mitochondrial cristae, we observed an increase of MIC27 (0.78 log_2_FC), MIC13 (1.1 log_2_FC), and MIC26 (1.4 log_2_FC) in Cyto-Mitos compared to Syncytio-Mitos. Additionally, Cyto-Mitos had a higher abundance of the inner membrane stomatin-like protein 2 (STOML2, 0.49 log_2_FC).

**Figure 3:**
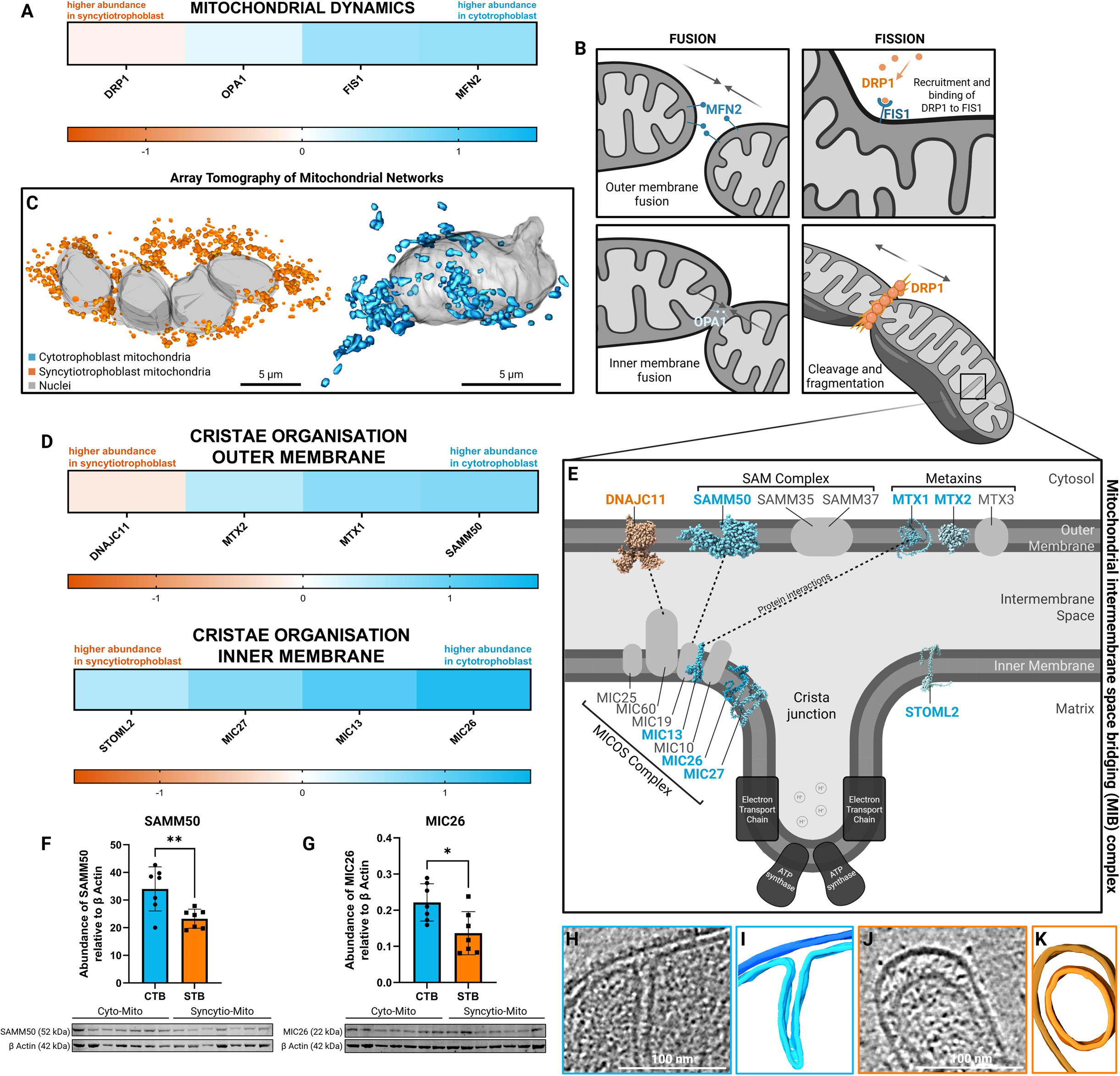
Proteomic analysis of Cyto- and Syncytio-Mitos to investigate the mechanisms involved in network fragmentation and structural remodelling. Heatmap graphs show the log_2_ fold change (log_2_FC) in protein abundance, with negative log_2_FC indicating a higher abundance in Syncytio-Mitos (orange) and positive log_2_FC indicating a higher abundance in Cyto-Mitos (blue). A) Abundance of proteins involved in mitochondrial dynamics and network structure, including DRP1, OPA1, FIS1, and MFN2. B) Diagrammatic representation of mitochondrial dynamics proteins and their role in network fusion and fission of the outer and inner membranes. C) Fusion of Cyto-Mitos (blue) and fragmentation of Syncytio-Mitos (orange) was confirmed by array tomography of placental villi. D) Abundance of proteins involved in the organisation of mitochondrial cristae, with the top row showing proteins embedded within the outer mitochondrial membrane, including DNAJC11, MTX2, MTX1, and SAMM50. The bottom row shows proteins embedded within the inner mitochondrial membrane, including STOML2, MIC27, MIC13, and MIC26. E) Diagrammatic representation of the mitochondrial intermembrane space bridging (MIB) complex, which includes protein complexes that are involved in organising mitochondrial cristae structure. Representative predicted protein structures have been shown, generated with Alphafold (Deepmind, UK).^75^ Western blotting was performed on mitochondrial isolates (n = 7 placentae) to validate the abundance of F) SAMM50 (52 kDa), and G) MIC26 (22 kDa), relative to β actin (42 kDa), between CTBs (blue) and the STB (orange) in a larger sample size. Grubb’s outliers tests were performed on western blotting data, followed by unpaired t-tests to examine significant variation, expressed as p < 0.05*, and p ≤ 0.01**. Representative blots are shown below the graphs. Images of the Cyto-Mito (H-I) and Syncytio-Mito (J-K) intermembrane spaces are shown, obtained from cryo-ET data.

To further validate our proteomic analysis, we performed western blotting and characterised two MIB proteins, one located within the outer (SAMM50) and the other residing within the inner mitochondrial membrane (MIC26), which had the greatest log_2_FC between trophoblast populations. These data showed changes consistent with our proteomic analysis, identifying a significantly greater abundance of SAMM50 and MIC26 in Cyto-Mitos compared to Syncytio-Mitos (Fig. 3F, G). In line with this, in our 3D reconstructions we observed a greater intermembrane space in Syncytio-Mitos compared to Cyto-Mitos, a feature that is governed by the aforementioned cristae organisation proteins (Fig. 3H-K).

### Subunit-specific variations in mitochondrial complexes between Cyto- and Syncytio-Mitos

To investigate the bidirectional relationship between cristae architecture and bioenergetic function, we assessed the composition of the ETC and ATP synthase. Our proteomic analysis identified distinct subunit-specific variations of complexes I-IV and ATP synthase, between Cyto- and Syncytio-Mitos. Specifically, we identified that complex I core subunits NDUFV1, NDUFV2, and NDUFS3 were more abundant in Cyto-Mitos, whereas NDUFS8, NDUFS2, and NDUFS7 were more abundant in Syncytio-Mitos. Our analysis also showed a higher abundance of nine accessory subunits in Cyto-Mitos, while seven accessory subunits were more abundant in Syncytio-Mitos (Fig. 4A). While there were no changes in SDHB of complex II in our dataset (Fig. 4B), there was a higher abundance of three complex III subunits in Cyto-Mitos, and three other subunits were higher in Syncytio-Mitos (Fig. 4C). Our analysis of complex IV determined that only COX4I1 was more abundant in Cyto-Mitos, whereas COX5B and COX5A were more abundant in Syncytio-Mitos (Fig. 4D). Investigation of ATP synthase subunits revealed that seven of the subunits were higher in Cyto-Mitos, while only ATP5O was higher in Syncytio-Mitos (Fig. 4E). To determine the position of each subunit identified in our proteomics analysis, subunit locations were mapped using representative protein structure models derived from HEK293 cells (Fig. 4F).^39,76–78^ Finally, to assess the capacity for ATP synthase dimerisation within Cyto- and Syncytio-Mitos, we confirmed by western blotting that Cyto-Mitos had a significantly higher abundance of dimerisation protein ATP5MK (Fig. 4G).

**Figure 4:**
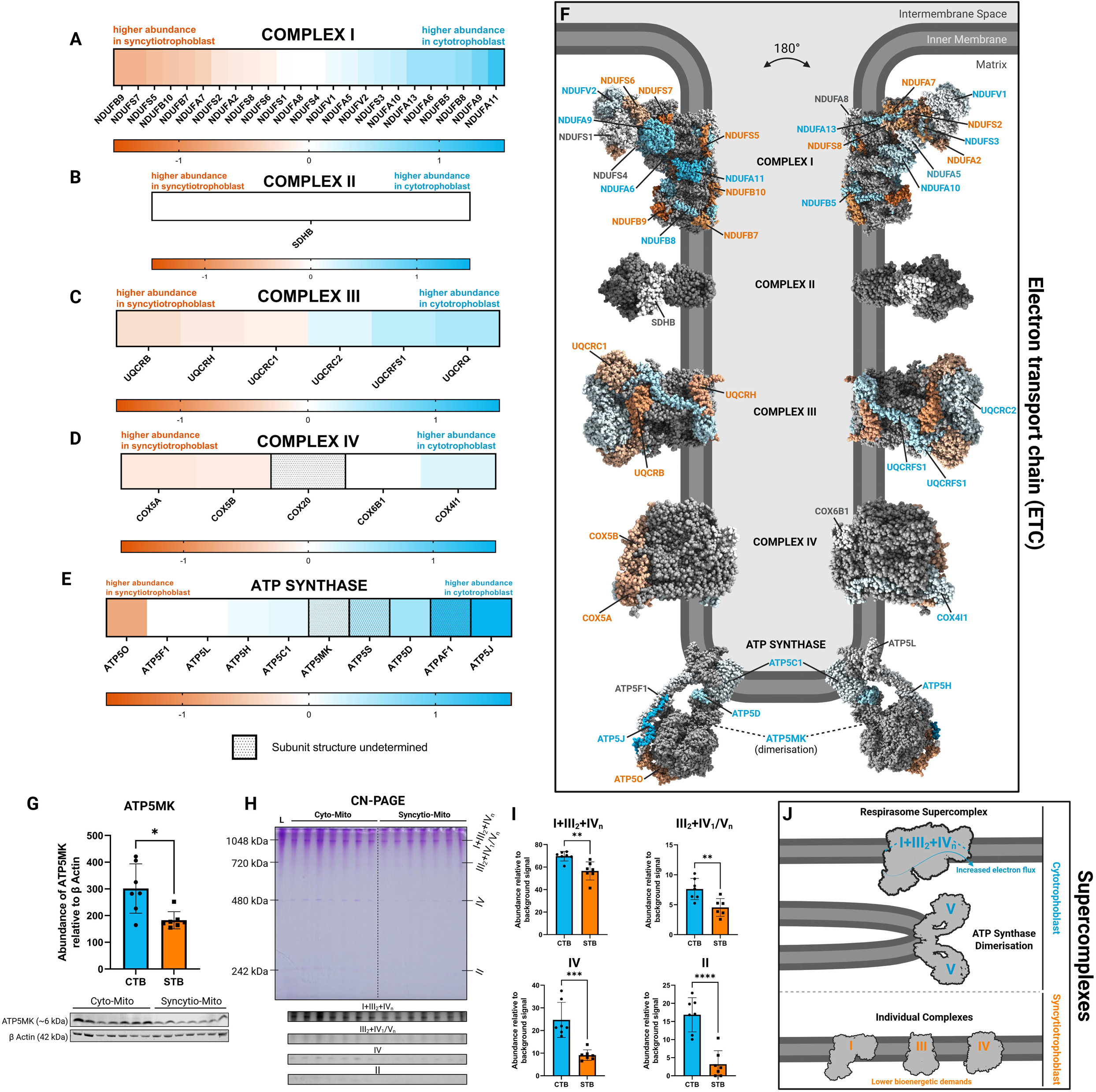
Proteomic analysis suggests altered ETC composition between the CTB and STB, to meet contrasting bioenergetic demands. Heatmap graphs show the log_2_FC in protein abundance, with positive log_2_FC indicating a higher abundance in the CTB (blue) and negative log_2_FC values indicating a higher abundance in the STB (orange). Graphs show the abundance of protein subunits of A) complex I, B) complex II, C) complex III, D) complex IV, and E) ATP synthase. Patterned heatmap squares (C and D) indicate subunits that have no solved structures in the representative models shown in F). F) Diagrammatic representation of the ETC and ATP synthase. As no protein models of placental mitochondria currently exist, representative cryo-EM models (obtained from HEK293F cells) have been used to show the location of each subunit, sourced from PDB (I: 5XTD, II: 8GS8, III: 5XTE, IV: 5Z62, V: 8H9V).^39,76–78^ G) Western blotting was performed using mitochondrial isolates (n = 7 placentae) to quantify the abundance of ATP5MK (7 kDa) relative to β actin (42 kDa), between CTBs (blue) and the STB (orange). H) CN-PAGE was performed using mitochondrial isolates (n = 7 placentae), and the representative membrane shows the abundances of supercomplexes I+III+IV (1048 kDa), III+IV, and individual ATP synthase (720 kDa), complex IV (480 kDa), and complex II (242 kDa). Quantitative analysis was performed on the excerpts of each band of proteins, to produce graphs in I). J) Summary diagram showing that Cyto-Mitos have elevated levels of respirasome supercomplexes and ATP synthase dimerisation, while Syncytio-Mitos maintain individual complexes. Grubb’s outliers tests were performed on western blotting data, followed by the Mann-Whitney U test to examine ATP5MK, and unpaired t-tests were used to examine CN-PAGE data. Significant variation is expressed as p < 0.05*, p ≤ 0.01**, p ≤ 0.001***, and p ≤ 0.0001****.

### Decreased supercomplex assembly in Syncytio-Mitos

Supercomplexes greatly increase kinetic efficiency and the capacity of mitochondria to synthesise ATP, compared to individual complexes. Clear native PAGE (CN-PAGE) was performed on placental mitochondrial isolates to determine differences in the abundance of supercomplexes between undifferentiated CTB and the differentiated STB. Cyto-Mitos had a significantly higher abundance of supercomplexes I+III+IV (p = 0.0025) and III+IV (p = 0.0065), compared to Syncytio-Mitos. Cyto-Mitos also had a significantly higher abundance of complex IV (p = 0.0006) and complex II (p < 0.0001)(Fig. 4H-J).

## Discussion

In this study, we conducted a multimodal 3D analysis of mitochondria from CTBs and the STB of the human placenta to study how changes in mitochondrial structure between differentiating cell types may translate to differences in function. Our imaging approach enabled both the investigation of mitochondrial networks in placental villous tissue in situ, and high-resolution analysis of inner membrane and cristae architecture in isolated human mitochondria. Using array tomography, we established that Cyto-Mitos have greater mitochondrial complexity and volume in comparison to the differentiated Syncytio-Mitos. Within individual mitochondria, cryo-ET revealed the internal architecture of Cyto- and Syncytio-Mitos, identifying for the first time that there are two unique subpopulations of mitochondria within progenitor cytotrophoblasts that have either lamellar or tubulo-vesicular cristae architecture. Notably, the architecture of both Cyto-Mito subpopulations was distinct from Syncytio-Mitos, which only consisted of globular cristae. In turn, our proteomic analysis of isolated mitochondria identified that Syncytio-Mitos have a reduced capacity to undergo fusion and organise internal cristae architecture, with a decrease in MICOS and SAM complexes compared to Cyto-Mitos. These results occur in conjunction with alterations in specific core and accessory subunits across complexes I, III, and IV of the ETC, and ATP synthase. This study is the first to characterise the presence of supercomplexes in the placenta and observe that Syncytio-Mitos have decreased abundances of supercomplexes I+III+IV and III+IV. Collectively, our findings contribute to the broader understanding of mitochondrial physiology by linking alterations in cristae organisation proteins, dimerisation, and supercomplex abundance, with 2D angularity measures. These measures are supported by a standardised visual metric that accounts for 3D topology, thereby facilitating the comparison of localised cristae curvature (curvedness).

### Mitochondrial dynamics

Our finding that Cyto-Mitos are larger and more elongated than Syncytio-Mitos, confirms prior 2D observations of the structural diversity of mitochondria in the placenta.^69^ Our novel and comprehensive 3D mapping enables a significantly more detailed analysis of the interconnected and complex network structures, demonstrating for the first time that Cyto-Mitos exhibit clustered and networked phenotypes. As mitochondrial networks are established via the process of fusion, this structural pattern of larger, elongated, and more extensive networks is associated with the higher abundance of fusion proteins MFN2 and OPA1.^79^ In contrast, the smaller Syncytio-Mitos that exist in more fragmented networks were associated with an increase in DRP1, which binds to FIS1 receptors to constrict and cleave mitochondria.^11,12^ Indeed, the change in mitochondrial morphology following CTB differentiation into the STB, aligns with previous studies that suggest the reduction of mitochondrial size in the STB supports steroidogenic roles.^90^ Specifically, steroidogenesis is facilitated by an increased mitochondrial surface area to volume ratio for the translocation of cholesterol to cytochrome P450, by which pregnenolone is synthesised.^90^ As first proposed, the remodelling of mitochondrial networks by fission may be critical for steroidogenic function, particularly since the human placenta lacks conventional steroidogenic acute regulatory proteins (StAR) that facilitate the translocation of cholesterol.^80–82^

In addition, we propose that fission may be necessary for the distribution of mitochondria throughout the multinucleated and functionally diverse STB, in a manner akin to neuronal axons.^14,83^ Furthermore, the presence of fragmented mitochondria in the STB is consistent with an increase in senescence-associated proteins in the STB such as P53, DCR2, and CDK inhibitors.^84,85^ Therefore, we suggest that in addition to functional specialisation, the fragmentation of the mitochondrial network is a process of quality control and preparation for mitochondrial clearing (mitophagy) in the aging STB.^8^ Our data also suggests a potential interplay between broader network dynamics and cristae architecture, observing reduced levels of STOML2 in Syncytio-Mitos. STOML2 plays an integral role in promoting inner membrane stability, and is suggested to restrict mitophagy, and promote mitochondrial biogenesis.^86,87^ Thus, reduced levels of STOML2 in Syncytio-Mitos suggests that there is greater mitochondrial clearance and impaired inner membrane stabilisation compared to Cyto-Mitos, and is supported by our observation of reduced fusion protein OPA1 in the STB.

### Regulation of internal architecture and cristae structure

The intermembrane space is critical for the accumulation of protons and the formation of an electrochemical gradient across the inner mitochondrial membrane, with MIB proteins manipulating cristae structure into proton-rich pockets.^29,74^ In this study, we have shown that cristae from Syncytio-Mitos do not exhibit the deep invaginated structure that we observed in Cyto-Mitos, and also have an increased intermembrane space. Mechanistically, this may be explained by the concurrent loss of MICOS and SAMM proteins in Syncytio-Mitos, which would ordinarily increase the capacity for inner membrane folding, tightening of cristae junctions, and the constriction of the intermembrane space to form funnel-like pockets.^29^ While prior studies have shown the association between MICOS and cristae architecture,^88^ our findings critically demonstrate for the first time that this interplay is linked to the remodelling of internal architecture and previously established changes in cellular demand.^67,70,71^ In conjunction with the aforementioned STOML2, Syncytio-Mitos also had reduced levels of MTX proteins which, although located within the outer mitochondrial membrane, are similarly critical for the stability of the cristae.^89–91^ In contrast, the high levels of MTX and STOML2 in Cyto-Mitos supports our observation of high cristae curvatures, which we and others attribute to a greater abundance of supercomplexes and ATP synthase dimers.^48,60,86^

Our ability to examine 3D cristae structure in the placenta for the first time, enabled us to identify two unique subpopulations of Cyto-Mitos, comprising either lamellar cristae or tubulo-vesicular cristae architecture. Contrastingly, only a homogeneous population of globular cristae is present in the differentiated Syncytio-Mitos. Our findings demonstrate that lamellar cristae of Cyto-Mitos have a higher surface area to volume ratio than the globular cristae of Syncytio-Mitos. While the surface area to volume ratio of tubulo-vesicular cristae did not significantly differ from that of globular cristae, the inclusion of the lamellar cristae significantly elevates the exchange efficiency and functional capacity of the broader Cyto-Mito population and, by extension, supports the higher bioenergetic demands of progenitor CTBs.^67,70,71,73^ Indeed, these cristae-specific observations highlight the importance of detailed sub-organelle structural analysis, contrasting the established dogma that we ourselves showed when imaging and assessing whole mitochondrial networks, in which Cyto-Mitos had a lower outer membrane surface area to volume ratio (Fig. 1). The inverse relationship we observed between the outer and inner membrane surface areas is not only a hallmark of the invaginated structure of mitochondrial cristae, but would also support the high bioenergetic capacity of Cyto-Mitos through the provision of additional surface area for the placement of the ETC.^67,70,71^ Moreover, the greater outer membrane surface area to volume ratio of Syncytio-Mitos can be attributed to the previously reported greater accumulation of P450, and the proposed prioritisation of steroidogenic functions,^69^ while the reduction in inner membrane surface area supports reduced cellular bioenergetic demands following trophoblast differentiation.^67,70,71^

To support and validate our 2D analysis of cristae angularity, we employed a standardised visual metric of curvedness that accounts for 3D topology and facilitates the comparison of localised cristae curvature. The acute angularity of lamellar apices and highly localised membrane curvedness we observed, further supports the greater bioenergetic capacity associated with Cyto-Mitos.^67,70,71^ Our findings are consistent with the theory that high curvature is induced by the presence of ATP synthase dimers, supercomplexes, OPA1, and MIB complexes, all observed in greater abundance within Cyto-Mitos.^28,29,46–48,74^ This demonstrates and validates the direct link between mitochondrial structure and function, whereby the angularity of cristae shapes the inner membrane to allow highly localised and concentrated electrochemical gradients at apices, thereby facilitating efficient proton funnelling through ATP synthase dimers.^48,49,51,92^ While the invaginated characteristics of lamellar cristae have previously been reported across species including yeast, green algae, and human skin fibroblasts,^48,51,92^ these studies do not measure 3D curvature, which accounts for complex membrane topology. Moreover, our assessment of tubulo-vesicular cristae identified uniformly distributed medium curvedness with pockets of angular ridges, which are absent in the globular cristae of Syncytio-Mitos. This finding is consistent with our previous statement pertaining to the abundance of ATP synthase dimers and supercomplexes in the Cyto-Mito populations.^46–48^ Globular cristae had the lowest curvedness, consistent with lower abundances of ATP synthase dimers and supercomplexes in Syncytio-Mitos and a decreased energetic capacity compared to Cyto-Mitos.^67,70,71^

While previous studies have compared local cristae curvature within individual mitochondria,^93–95^ our approach enables the normalised comparison of multiple cristae from different mitochondrial populations. In addition to the two distinct Cyto-Mito subpopulations described, we observed a mitochondrion consisting of a combination of both lamellar and tubulo-vesicular cristae architectures (Fig. S2). The co-existence of multiple cristae architectures within a singular mitochondrion has been suggested to enable a greater diversity of function in a variety of single-cell eukaryotes.^26^ Although singular in our dataset, the collective presence of at least two concurrent subpopulations of lamellar and tubulo-vesicular Cyto-Mitos poses the exciting prospect that we observed the process of “cristae differentiation”. As such, the development of mitochondrial subpopulations may precede, or accompany, the transition of proliferative CTBs to the previously identified pre-fusion CTBs in preparation for differentiation into STB.^96^ Our findings are consistent with the observations of Ryu et al.,^97^ who recently identified metabolically distinct mitochondrial subpopulations within mouse embryonic fibroblasts. Future work confirming the presence of mitochondrial subpopulations and distinct cristae architectures in situ, would require the application of cryo-ET to study vitrified, thin regions of tissue prepared by a plasma focused-ion-beam scanning electron microscope (plasma FIB-SEM).^98^

### The interplay between cristae structure and function

The transformation of cristae structure following trophoblast differentiation, and the subsequent loss of inner membrane surface area for mitochondrial functions of ATP synthesis in Syncytio-Mitos, supports previously reported lower cellular demands for ATP in the STB.^71^ In addition, we characterised the ETC, revealing distinct variations in the composition of complexes between Cyto- and Syncytio-Mitos. While the abundance of core subunits within Complex I was equally varied between the populations, we found that the majority of accessory subunits were more abundant in Cyto-Mitos. Since core subunits are necessary for electron transport, and remain unchanged between the populations, the loss of accessory subunits in Syncytio-Mitos suggests that there is a reduced dependence upon complex I structure, consistent with previous examinations of complex I and II linked respiration.^67^ Moreover, reductions in complex I accessory subunits have previously been shown to significantly impair the bioenergetic capacity of mitochondria within human embryonic kidney cells.^32^ Through a similar mechanism, the reduction of complex I accessory subunits in Syncytio-Mitos may be critical for the shift in cellular requirements following the differentiation of CTBs into the STB.^32^ Although this study observed no change in complex II subunits, it has previously been identified that there is a lower abundance of SDHA in Syncytio-Mitos.^71^ Subunits were mostly equally abundant in complexes III and IV.

The majority of ATP synthase subunits were higher in Cyto-Mitos. This finding is consistent with previously published results regarding ATP synthase subunits ATP5α and ATP5β.^71^ As the most critical stage for ATP synthesis, the increased abundance of ATP synthase subunits may underpin the higher bioenergetic capacity and respiratory rate of Cyto-Mitos.^67,70,71^ In addition, we observed a higher abundance of ATP5MK in Cyto-Mitos (confirmed by western blotting), which suggests a greater propensity for ATP synthase dimerisation in the CTB, a feature conserved across highly efficient bioenergetic mitochondria from yeast, bovine and human cells.^46,78,99^ Further, the increase in ATP5MK may contribute to the greater membrane curvature observed in Cyto-Mitos, with the assembly of ATP synthase dimers shown to induce membrane curvature, and localisation of electrochemical gradients at cristae apices.^48,50,95^ While not directly assessing dimerisation, our findings are consistent with the high curvedness of lamellar cristae apices, and acute angular pockets observed in tubulo-vesicular cristae in Cyto-Mitos relative to the globular cristae of Syncytio-Mitos.

### The role of supercomplexes in shaping cristae architecture

In addition to ATP synthase dimers inducing curvature, cristae architecture has been proposed to be bidirectionally influenced by supercomplex assembly. It has previously been suggested that increased cristae curvature promotes the assembly of supercomplexes, through OPA1-linked mechanisms that contribute to the organisation of internal architecture.^100^ A concept supported by our observation of increased OPA1 in Cyto-Mitos, in conjunction with greater membrane curvedness at cristae apices. Moreover, research has established that the placement of individual complexes may cause small deviations to the inner mitochondrial membrane, while rows of supercomplexes have been shown to significantly bend the inner membrane by up to 130°.^60^ The increased abundance of supercomplexes in Cyto-Mitos may explain the high curvedness in lamellar and tubulo-vesicular cristae within Cyto-Mitos, relative to Syncytio-Mitos. Moreover, the unexplained variations in ETC composition, particularly in accessory subunits, may promote greater assembly of the supercomplexes I+III+IV and III+IV in Cyto-Mitos. The subunit-specific variations we identified may also support the co-existence of supercomplexes and individual complexes in a dynamic state of plasticity.^55^ While supercomplexes in Cyto-Mitos may support increased bioenergetic demands and higher respiratory rates, reduced assembly in Syncytio-Mitos may explain previously reported lower demands for energy in the STB, and a shift in functionality from ATP synthesis towards steroidogenesis.^69,71^ In addition, we propose that the remodelling of mitochondrial structure following trophoblast differentiation is necessary to reduce the reliance of Syncytio-Mitos upon oxidative phosphorylation. This, in turn, limits the overutilisation of oxygen and substrates by Syncytio-Mitos, which would otherwise diffuse through the STB into fetal vessels, and support fetal development.

### Limitations

In this study, we showcased the advantage of using a multimodal imaging approach, exploiting the benefits of both array tomography and cryo-ET to study the diversity of mitochondrial structure in the placenta. While array tomography has the capacity to image whole and intact mitochondrial networks of the placenta, the technique is unable to be applied to study internal architecture and cristae structure due to resolution limits. This limitation is resolved by the incorporation of cryo-ET, which can acquire images with greater resolution and magnification. The mitochondrial isolation procedure used in our cryo-ET imaging has previously been validated to maintain mitochondrial capacity and respiratory function.^67^ Although our electron microscopy analysis was limited by the number of patient samples (n = 2 placentae for array tomography and n = 2 placentae for cryo-ET), we increased the number of mitochondria analysed, making statistical comparisons between mitochondria of the two cell populations (n = 354 mitochondria for array tomography and n = 16 mitochondria for cryo-ET), rather than assessing interpatient differences that would be used to investigate pathological instead of healthy mitochondrial structure. To support our 2D analysis of mitochondrial cristae, this study employs a workflow for calculating the gross 3D topology and curvature (curvedness). Curvature estimation is notably challenging due to the interference of segmentation and 3D reconstruction artefacts, and previously published examples focus on comparing local curvature within a singular mitochondrion.^94,95^ However, our approach of membrane reconstruction and smoothing enabled the characterisation of functionally pertinent gross membrane curvature, and the comparison of multiple cristae architectures from different mitochondrial populations. To support our imaging observations, we conducted an in silico analysis within a previously published shotgun proteomics dataset,^71^ in which we compare protein abundance across specific pathways including MIB and ETC complexes. This enabled the comparison of log_2_FC, which provides important insights into differences between Cyto and STB-Mitos. While statistical analysis of these differences was not performed in this study, we validated and confirmed the major findings from the proteomics data by performing western blotting on an additional seven placentae. CN-PAGE was performed to assess supercomplex abundance, however, this technique has a lower separation resolution than the alternative blue native PAGE (BN-PAGE) technique. However, we identified that the addition of Coomassie dye (a requirement of BN-PAGE) interfered with the separation of Cyto- and Syncytio-Mito native complexes – a notable advantage of CN-PAGE.^101^

### Conclusion

The placenta is a unique model for studying diverse mitochondrial structures within a single, bioenergetically dynamic organ. This study has revealed the importance of assessing 3D topology when considering the mechanisms that govern mitochondrial structure and function as cells differentiate. Specifically, we have shown that progenitor CTBs adopt a more complex network of large, elongated, and branched mitochondria, which transition into fragmented, spherical, and compact mitochondria that are distributed across the multinucleated STB. The remodelling of mitochondrial structure is consistent with the reduction of fusion proteins MFN2 and OPA1, and an increase in fission protein DRP1 in the STB. We identified a complex interplay between gross morphology and cristae architecture, governed by MIB constituents including the MICOS complex and STOML2 of the inner mitochondrial membrane, and the SAM and MTX complexes of the outer mitochondrial membrane. The reduction of cristae organisation proteins in Syncytio-Mitos, supports our cryo-ET observations of isolated mitochondria, in which we observed the absence of invaginated, lamellar cristae, and the formation of globular cristae in Syncytio-Mitos. In addition, our study assessed the complex topology and 3D curvature of lamellar apices, and the localisation of curvature pockets in tubulo-vesicular cristae, found exclusively within the second subpopulation of Cyto-Mitos. Our findings suggest that these dramatic differences in morphology and cristae structure are in part, influenced by alterations to core and accessory subunits of the ETC, impacting the ability of ATP synthase to form dimers, and the assembly of supercomplexes I+III+IV and III+IV, both of which were more abundant in Cyto-Mitos. In summary, this study demonstrates the advantages of adopting a multimodal approach of array tomography and cryo-ET imaging in combination with proteomic analysis, to study diverse mitochondria and the mechanisms that link structure and function.

## Methods

### Ethical Approval

Ethical approval was granted by the University of Newcastle Research Ethics Committee (H-382-0602), Hunter New England Health Human Research Ethics Committee (02/06/12/.13), Mercy Health Human Research Ethics Committee (R11/34), Griffith University Human Research Ethics Committee (MSC/05/15HREC), Queensland Government Human Research Ethics Committee (HREC/14/QPCH/246), and Site Specific Assessment (SSA/15/HNE/291) in compliance with the Declaration of Helsinki standards. With informed consent, placental samples were collected from healthy full-term pregnancies (38-40 weeks) at the John Hunter Hospital (NSW, Australia) and Mercy Hospital for Women (VIC, Australia). All reagents were sourced from Sigma Aldrich, Australia, unless otherwise specified.

### Array Tomography

#### Array Tomography Tissue Preparation

Placental villous tissue was collected using a 6 mm biopsy punch, from placentae of two healthy full-term pregnancies. Tissue samples were first washed in ice-cold PBS, then fixed in a solution of 1% (w/v) glutaraldehyde, 1.5% paraformaldehyde, 5 mM calcium chloride in 0.1 M sodium cacodylate. Samples were then prepared using the previously published reduced osmium fixation (ROTO) technique^102^ and then embedded in Procure 812 resin (ProSciTech, Australia).

#### Array Tomography Image Acquisition and Processing

Resin-embedded placental tissue was serially sectioned at a thickness of 80 nm, using a Leica UC7 ultramicrotome (Leica Biosystems, Germany), and recovered onto silicon wafers that were sputter coated with 10 nm of carbon. Sections were then imaged using a FEI Teneo VolumeScope scanning electron microscope (Thermo Fisher Scientific, USA), with a voltage of 3 kV and current of 0.2 nA, acquired using MAPS software (Thermo Fisher Scientific, USA). Original images had resolution of either 6 or 8 nm, and were subsequently binned to 16 nm or 24 nm respectively, to improve the signal to noise ratio. Image series were then realigned using the IMOD package ETomo interface (University of Colorado, USA).^103^

#### Array Tomography Image Analysis and Segmentation

Image series were imported into the IMOD software suite (University of Colorado, USA), and manual segmentation was performed on CTB membranes, nuclei, and arbitrary boundaries of STB of approximately equivalent volume to the CTB – due to the continuous nature of the STB. CTBs were identified based on their morphology and proximity to the STB, while the STB was identified by its distinct multinucleated morphology and microvilli.^66,72,104^ Mitochondria were identified for segmentation based on the electron density resulting from the double membrane. To investigate and quantify the structure of mitochondrial networks, all mitochondria within each cell type were initially segmented as a single object mesh. Subsequently, to characterise volumetric properties of each mitochondrion, four randomly selected Z-slices per cell type were selected, and all mitochondria within each selected slice were manually segmented as new individual object meshes (henceforth referred to as the ‘subset’). Mitochondria with obscured views or partial volumes were excluded from analyses. Each mitochondrion of the subset was meshed to form a 3D reconstruction, resulting in a subset of Cyto-Mitos (n = 113) and Syncytio-Mitos (n = 241). Elongation in the XY plane was calculated as a ratio from manual 2D measurements of maximum length and width. The sphericity (*ψ*) of each mitochondrion was calculated using the published formula^105,106^

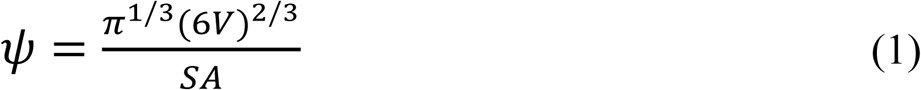

where *V* is volume and *SA* is surface area.

Additionally, we applied the previously published mitochondrial complexity index (MCI) formula,^107^ to quantify the structural complexity of each mitochondrion, as a method of investigating the degree of network interconnectedness and mitochondrial fusion occurring with neighbouring mitochondria

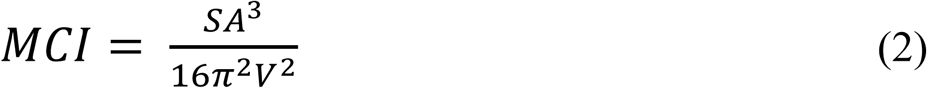

as previously described.^107^ While sphericity asymptotically reaches a value of one for a perfect sphere, MCI does not have theoretical limits – allowing the examination of more complex and clustered mitochondrial structures.^107^

### Cryo-electron Tomography (cryo-ET)

#### Liquid Ethane Vitrification

Mitochondria were freshly isolated from two healthy full-term placentae via sequential differential centrifugation as per published protocols.^67^ From each placenta, three tissue biopsies were used to produce pooled and concentrated Cyto- and Syncytio-Mito isolates. As soon as possible after birth, isolated mitochondrial samples were resuspended in 50 µL of isolation buffer (250 mM sucrose, 0.5 mM sodium EDTA, 10 mM tris) and vitrified in liquid ethane using the Vitrobot Mark IV System (Thermo Fisher Scientific, USA). To do this, 3 µL of each sample was blotted for 5 seconds, with a blot force of 7, onto glow discharged Quantifoil R 1.2/1.3 300 mesh Au grids (Quantifoil, Germany). Glow discharging was performed using a Pelco EasiGlow (Pelco, USA) with a plasma current of 15 mA for 30 seconds, ultimate pressure set to 0.30 mbar, and stable pressure of 0.39 mbar.

### Cryo-ET Image Acquisition

Vitrified grids were clipped into AutoGrids, and imaged using the FEI Titan Krios 300 keV field emission gun TEM (Thermo Fisher Scientific, USA), equipped with a Gatan BioQuantum energy filter, and K3 camera system (AMETEK, Inc., USA). Single axis bidirectional tilt series were acquired using Tomography Software (Thermo Fisher Scientific, USA), with a range of ± 50°, at 3° increments and 3 second exposures for Cyto-Mitos, or 2° increments and 2 second exposures for Syncytio-Mitos. We identified that a higher exposure was necessary to improve the visualisation of inner membranes within the large and electron-dense Cyto-Mitos, whereas the smaller Syncytio-Mito inner membranes were less obscured. The compromise, however, was that the incremental step size was also increased to limit the total electron dose and prevent beam-induced damage.^108^ Both sample types were imaged at 42,000 x magnification, with a defocus range between -5 µm and - 8 µm. Zero-loss imaging was performed with an energy filter width set to 15 eV, using the counted and super resolution modes on the K3 direct detector, and subsequently binned by 2 for reconstruction. The maximum accumulated electron dose for tomograms was 100 e^-^/Å^2^, with pixel sizes ranging between 2.1 Å and 3.4 Å.

### Cryo-ET Tomogram Reconstruction

Tilt series were processed using the IMOD software suite (University of Colorado, USA).^103^ Exposures with significant beam shift or obstructed views were excluded from the reconstruction process. Tilt series were coarsely aligned using Tiltxcorr and finely aligned using fiducial-less patch tracking, prior to 2 x binning to improve the signal to noise ratio. Tomograms were reconstructed using the back projection method, with a SIRT-like filter set to 20 iterations, using the Hamming-like filter.^109^

### Cryo-ET Segmentation

Reconstructed tomograms were imported into AMIRA (Thermo Fisher Scientific, USA), and denoised using the median and non-local means filters to improve the contrast of mitochondrial membranes.^110,111^ Manual segmentation of Cyto-Mitos and Syncytio-Mitos (n = 8 mitochondria per cell type) was performed every 5-10 Z-slices and then interpolated – with interpolated segmentations checked for inaccuracies and significant deviations from the true membrane line. Slices with poor visualisation of deeper internal membranes were excluded from segmentation. Segmentations were meshed to produce 3D reconstructions of mitochondrial outer membranes and cristae.

### Quantification of Intermembrane Distance

For each mitochondrion, the tomographic slice with the best membrane visibility was chosen for all 2D measurements. Each mitochondrion was split into four cartesian quadrants (Fig. S1), and for each quadrant, the perpendicular distance between the outer and inner membrane was measured if it presented with clearly visible membranes (resulting in up to four intermembrane distance measurements per mitochondrion).

### Quantification of Cristae Length and Width

The maximum XY length of each crista was measured, followed by the perpendicular maximum XY width. Elongation was calculated as a ratio between the length and width measurements (Fig. S1).

### Quantification of Cristae Angle

For each lamellar cristae, the XY angles of junctions and apices were measured, while the cristae angle for tubulo-vesicular and globular cristae was measured using the quadrant system, with at least one angle measurement taken from each quadrant. Additional angles were measured for cristae with multiple deviations in membrane curvature (Fig. S1).

### Quantification of Surface Area and Volume

Cristae surface area was calculated using the membrane segmentations, whereas cristae volume was calculated using segmentations with filled cristae lumen. It was necessary to develop two distinct reconstructions to ensure that surface area measurements were not skewed by the flat ‘caps’ that were required to enclose the lumen of cristae for volume calculations. The volume of lamellar cristae was calculated by connecting the cristae junctions with a faux inner boundary membrane, thus enclosing the lumen, similar to a previously described workflow.^94^ Volume data was normalised to the number of tomographic slices for each mitochondrion.

### Quantification of 3D Curvature

To characterise the complex topology and 3D curvature of cristae structure, six cristae reconstructions were first imported into MeshLab (ISTI-CNR, Italy), cleaned to remove flat “caps” that enclosed the cristae, and the Laplacian smoothing filter (set to 500 iterations) was applied to mitigate the interference of segmentation artefacts upon curvature estimation, while preserving large-scale features and gross topology.^112^ Smoothing was consistently applied to all reconstructions prior to the analysis of curvature, ensuring that segmentation edge effects were minimised. Smoothed reconstructions were then imported into ParaView (Sandia National Laboratories, USA), and principal curvatures were derived from mean and gaussian curvature calculations using the python calculator tool. Principal curvatures were subsequently used to calculate curvedness

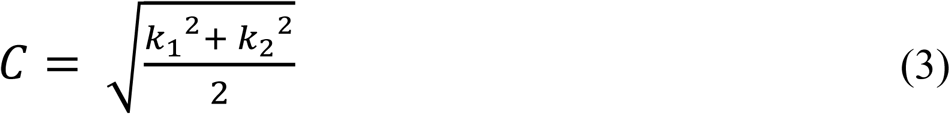

where *C* is curvedness, *k*_1_ is maximum curvature, and *k*_2_ is minimum curvature.^113^

As previously published, curvedness assesses the magnitude of curvature irrespective of direction, and is therefore more appropriate for comparing complex cristae topologies compared to mean and gaussian curvature measurements.^113^ The notable advantage of using ParaView to calculate curvedness, was the normalisation of colour maps, enabling a standardised comparison of multiple cristae reconstructions from different tomographic data.

#### Statistical Analysis of Electron Microscopy Data

Array tomography and cryo-ET quantification data was processed using GraphPad Prism (GraphPad Software, USA), and first examined for outliers using Grubb’s Outliers test. The Shapiro-Wilk test was performed on all groups to assess the normality of distribution. All array tomography data, and intermembrane distance was examined using the Mann-Whitney U test. The Kruskal-Wallis and Dunn’s multiple comparisons tests (post-hoc) were applied to compare all cryo-ET metrics between lamellar, tubulo-vesicular, and globular cristae architectures, except for volume – which was normally distributed and therefore analysed using the Brown-Forsythe and Welch ANOVA test (post-hoc). Significant variation between groups was established as p < 0.05.

### In Silico Analysis of Proteomic Data

This study performed an in silico analysis of previously acquired LC-MS data.^71^ As previously described, mitochondria were isolated from placental villous tissue of healthy full-term pregnancies,^67^ and LC-MS was performed using enriched fractions of Cyto- and Syncytio-Mitos (n = 3 placentae).^71^ In contrast to the previous publication which only presented proteins deemed significant following Fisher’s exact test independent to fold change, this study performed an in silico analysis of the dataset to assess protein changes between mitochondrial populations within specific pathways. To determine the pathways, the following terms were used; NADH dehydrogenase [ubiquinone] (n = 24 subunits), succinate dehydrogenase [ubiquinone] (n = 1 subunit), cytochrome b-c1 (n = 6 subunits), cytochrome c oxidase (n = 5 subunits), ATP synthase (n = 10 subunits), in addition to select proteins that reside within mitochondrial dynamics (n = 4 proteins), or MIB complexes (n = 8 proteins). Proteins were normalised to Cyto-Mitos as previously described,^71^ and the log_2_ fold change (log_2_FC) identifies directionality and relative abundance between the populations. Proteins with less than three unique peptide sequences identified by LC-MS were excluded from analysis to improve the reliability and reproducibility of our results. Heatmap graphs were produced using GraphPad Prism (GraphPad Software, USA), with a positive log_2_FC representing a greater abundance in Cyto-Mitos, and a negative log_2_FC as a greater abundance in Syncytio-Mitos.

### Immunoblotting

#### Clear-native PAGE (CN-PAGE)

To perform CN-PAGE, Cyto- and Syncytio-Mito isolates (n = 7 placentae) were resuspended in aminocaproic acid buffer (1.5 M aminocaproic acid, 50 mM Bis-Tris) and the protein concentration measured using a BCA assay kit. As previously described,^114^ digitonin was added to achieve a 1:1 protein/detergent ratio for each sample, and incubated for 10 min on ice. Following incubation, samples were centrifuged at 20,000 x g for 30 min at 4° C and the pellet was discarded. To the supernatant, 1 x native loading buffer (aminocaproic acid, 1 M Bis-Tris, 500 mM EDTA, Coomassie Brilliant blue G 250) was added, and 10 µL of each sample (5 µg of protein per well) was loaded into the wells of a 4-12% Bis-Tris gel (Invitrogen, USA). Protein size was assessed using 3 µL of NativeMark Unstained Protein Standard. Following this, the cathode and anode chambers of the electrophoresis tank were filled with 50 mM Bis-Tris. Electrophoresis was performed at 4 °C to prevent overheating, at a constant voltage of 300 V, until the lowest band of proteins reached the bottom of the gel. After washing the gel in 50 mM Bis-Tris for 30 minutes, imaging was performed using the Amersham Imager 600 (GE HealthCare, USA).

#### Western Blotting

Western blotting was performed as previously described.^71^ Cyto- and Syncytio-Mito isolates (n = 7 placentae) were resuspended in 250 µL of RIPA buffer (100 mM Tris, 300 mM sodium chloride, 10% Nonidet P40, 10% sodium deoxycholate, 1% sodium dodecyl sulfate, 5 x Complete Mini Protease Inhibitor Cocktail tablet (Roche Diagnostics, Australia), pH 7.4). Sample lysates were prepared to a normalised concentration of 40 µg/µL, with Milli-Q H_2_O, NuPAGE Sample Reducing Agent, and NuPAGE Loading Dye (Thermo Fisher Scientific, USA). Lysates were heated to 90 °C for 10 minutes, then stored at -80 °C until use. Electrophoresis was performed with chilled 1 x NuPAGE MOPS SDS buffer (Invitrogen, USA), for 10 minutes at 100 V and then 90 minutes at 200 V. Transfer was performed using 1 x transfer buffer (200 mL 10 x transfer buffer (72.5 g tris hydrochloride, 36.6 g glycine, 1 L H_2_O), 400 mL methanol, 1400 mL H_2_O), for 10 minutes at 100 V and 60 minutes at 200 V. Membranes were washed in 5 mL PBST wash buffer (100 mL 10 x PBS, 2% Tween 20, 900 mL H_2_O) for 30 minutes at 4 °C, and then blocked for 1 hour using 5 mL of Odyssey Blocking Buffer (LI-COR Biosciences, USA). Following a 15 minute PBST wash, membranes were incubated with the following primary antibodies at a concentration of 1:250; anti-apolipoprotein O or MIC26 (ab246865, Abcam, UK) anti-SAMM50 (ab133709, Abcam, UK), anti-beta actin (ab8226, Abcam, UK), and anti-ATP5MK (pa569671, Thermo Fisher Scientific, USA), overnight at 4 °C. Following a 30 minute PBST wash, membranes were incubated in 1:20,000 IRDye 680LT goat anti-rabbit or IRDye 800CW goat anti-mouse secondary antibody (LI-COR Biosciences, USA) for 1 hour. After a final 15 minute PBST wash, membranes were imaged using the Odyssey M Imager (LI-COR Biosciences, USA).

#### Analysis of Membrane Blots

CN-PAGE and western blotting membrane images were analysed using Image Studio (LI-COR Biosciences, USA), and blot signal intensities were quantified relative to background noise. Supercomplexes were identified by molecular weight, as previously published.^115^ Western blots were run in experimental duplicates, with both membrane quantifications averaged prior to statistical analysis. GraphPad Prism (GraphPad Software, USA) was used to perform Grubb’s Outliers Test, followed by the Mann-Whitney U test to assess ATP5MK, and unpaired t-tests for all other proteins. Significant variation was established as p < 0.05.

## Acknowledgements

This study was supported by funding received from the John Hunter Hospital Charitable Trust, Newcastle Permanent Emerging Innovator Award, Hunter Medical Research Institute, and the National Health & Medical Research Council (NHMRC GNT2026065) Investigator Grant, and the Australian Government Research Training Program (RTP) Scholarship. We wish to thank and acknowledge the use of the University of Wollongong Cryogenic Electron Microscopy Facility at Molecular Horizons, under the management of Dr. James C. Bouwer and leadership of Executive Dean, Faculty of Science, Medicine and Health Senior Professor Eileen McLaughlin. We also acknowledge the electron microscopy facilities and staff of the Ian Holmes Imaging Centre, Bio21 Molecular Science and Biotechnology Institute and the University of Melbourne. We would like to express our gratitude to the research midwives and clinical staff of the John Hunter Hospital (Newcastle, Australia), and the Mercy Hospital for Women (Melbourne, Australia), and to the patients who donated placentae for this research.

