## Supplementary Figures for "Electron tomography reveals mitochondrial network and cristae remodelling during cell differentiation in the human placenta"

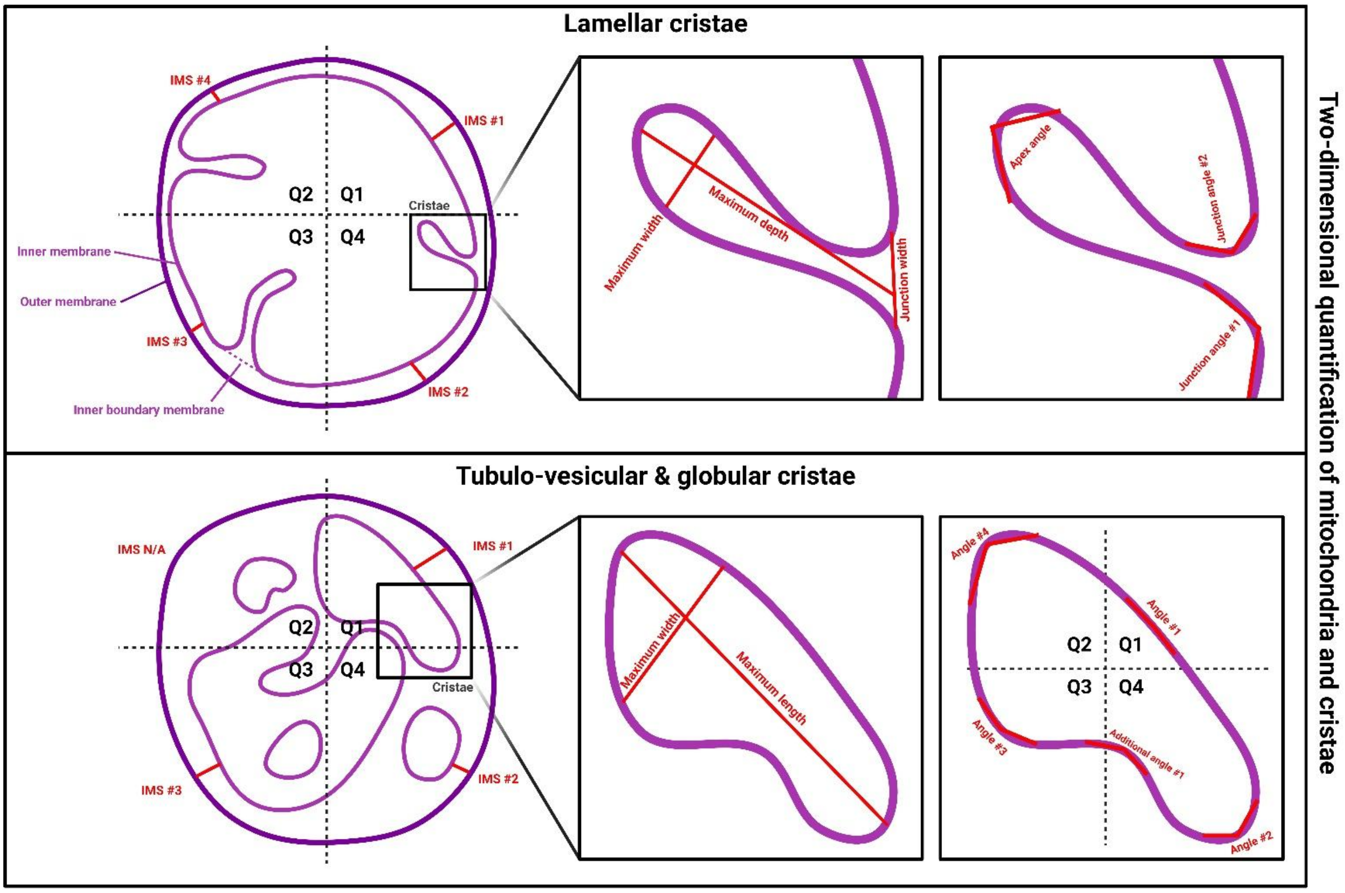


**Figure S1: Quantification of 2D mitochondrial and cristae structure.** Representative diagram showing the method for acquiring 2D metrics for lamellar cristae (above) or tubulo-vesicular/globular cristae (below). The tomographic slice with the best presentation of clear membranes was chosen for performing measurements. Each mitochondrion was split into four cartesian quadrants, and an intermembrane space measurement was made in each quadrant if there were clear parallel outer and inner membranes. For each lamellar cristae, the maximum depth and perpendicular maximum width was measured. Each apex and opposing junction angles were measured for all lamellar cristae. For tubulo-vesicular and globular cristae, maximum length and perpendicular maximum width was measured, and the cristae cross-section was split into four cartesian quadrants to measure the membrane curvature in each quadrant. In addition, angle measurements were recorded for any significant deviatoric curvatures (as shown by the representative “additional angle #1”).

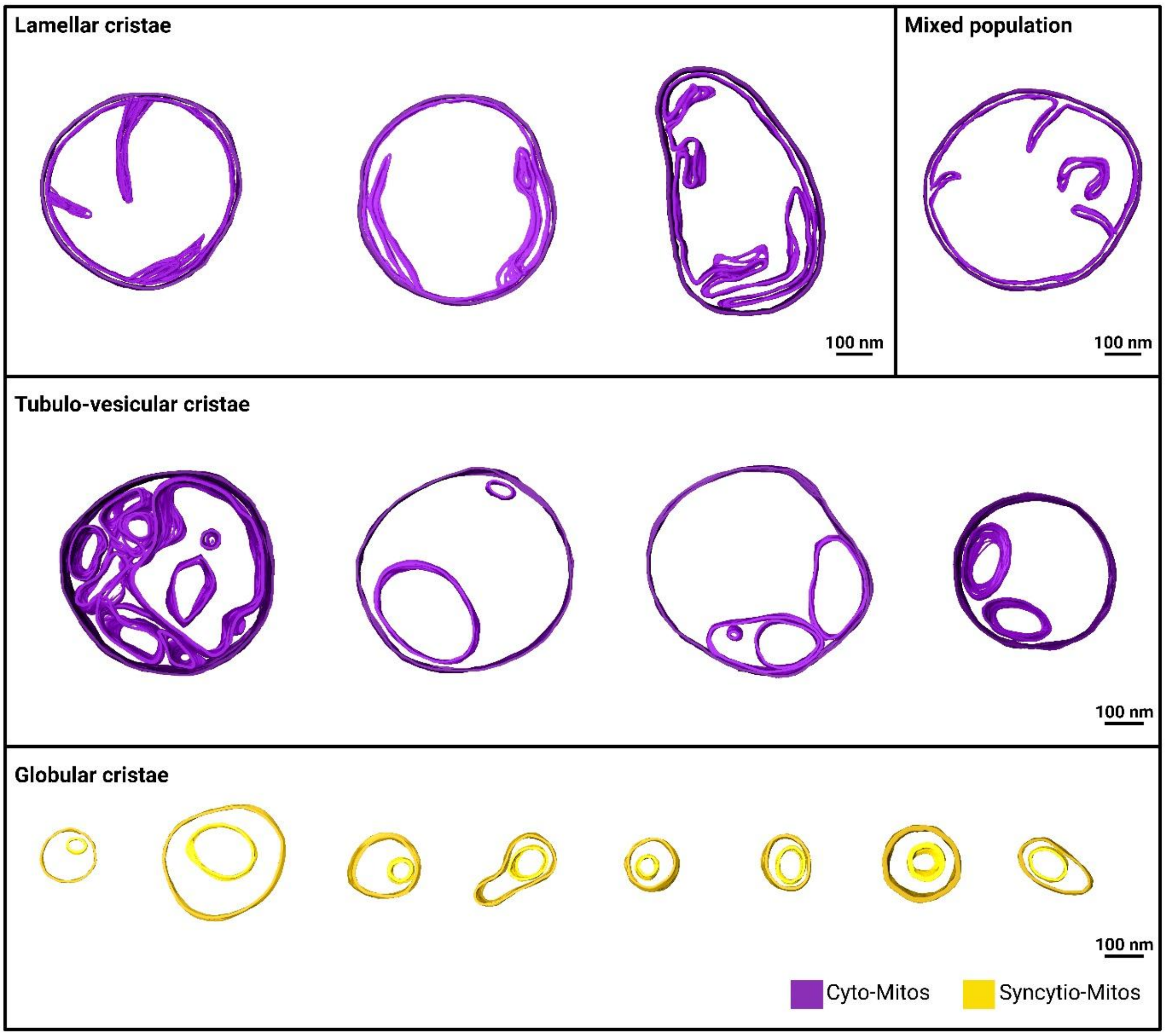


**Figure S2: Overview of all mitochondria analysed by cryo-ET.** 3D reconstructions were produced from tomographic data of Cyto- and Syncytio-Mitos. Two primary mitochondrial subpopulations are observed in the CTB (purple), consisting of lamellar cristae or tubulo-vesicular cristae, in addition to what may be a third subpopulation consisting of a combination of both cristae architectures “mixed population”. A homogeneous mitochondrial subpopulation is observed in the STB (yellow), of globular cristae – with the exception of one mitochondrion that consisted of irregular shaped cristae (dashed box). Scale bar represents 100 nm.
